# InterLabelGO+: Unraveling label correlations in protein function prediction

**DOI:** 10.1101/2024.06.26.600816

**Authors:** Quancheng Liu, Chengxin Zhang, Lydia Freddolino

## Abstract

**Motivation:** Accurate protein function prediction is crucial for understanding biological processes and advancing biomedical research. However, the rapid growth of protein sequences far outpaces the experimental characterization of their functions, necessitating the development of automated computational methods.

**Results:** We present InterLabelGO+, a hybrid approach that integrates a deep learning-based method with an alignment-based method for improved protein function prediction. InterLabelGO+ incorporates a novel loss function that addresses label dependency and imbalance and further enhances performance through dynamic weighting of the alignment-based component. A preliminary version of InterLabelGO+ achieved a strong performance in the CAFA5 challenge, ranking 6th out of 1,625 participating teams. Comprehensive evaluations on large-scale protein function prediction tasks demonstrate InterLabelGO+’s ability to accurately predict Gene Ontology terms across various functional categories and evaluation metrics.

**Availability and Implementation:** The source code and datasets for InterLabelGO+ are freely available on GitHub at https://github.com/QuanEvans/InterLabelGO. The software is implemented in Python and PyTorch, and is supported on Linux and macOS.

**Contact:** (LF) and (CZ)

## 1. INTRODUCTION

With rapid advancements in high-throughput sequencing technologies, the space of known protein sequences has expanded at an unprecedented rate. Despite this surge in sequence data, our ability to experimentally determine protein functions remains significantly limited. This discrepancy has led to a growing divide between the vast number of identified sequences and the relatively small subset with experimentally confirmed functions [5]. To address this challenge, it is crucial to develop automated computational methods capable of accurately predicting protein functions on a large scale.

Many computational approaches have been developed to address the protein function prediction problem. Sequence template-based methods, such as GOtcha [11] and Blast2GO [6], rely on sequence similarity to transfer functional annotations from well-characterized proteins to query sequences. Structural template-based methods, such as COFACTOR [17] and ProFunc [9], utilize structural information to infer protein functions. However, these methods have limitations in capturing distant evolutionary relationships and struggle with proteins that lack close homologs with known functions. Literature-based methods, such as DeepText2GO [14], aim to extract functional information from the abstracts of protein-related publications using natural language processing techniques. These methods can capture valuable functional insights from unstructured text data but face challenges in terms of scalability, information retrieval accuracy, and the availability of relevant literature for all proteins. Deep learning-based methods, such as DeepGOPlus [7] and TALE [2], extract sequence embeddings using convolutional neural networks (CNN) and transformers, respectively, to predict protein functions. While these methods can capture complex patterns in protein sequences, they may not fully exploit the rich evolutionary information contained in protein language models.

To overcome the limitations of existing methods and address the large-scale and multi-label nature of protein function prediction, we developed InterLabelGO+, a hybrid approach that integrates InterLabelGO, a deep learning-based method leveraging protein language models (PLM), with AlignmentKNN, an alignment-based method, for improved performance. InterLabelGO+ incorporates several key features to tackle the challenges of protein function prediction. First, InterLabelGO introduces a novel loss function that integrates label dependency and addresses label imbalances, enabling the model to capture complex functional relationships and handle the skewed distribution of functional annotations. Second, InterLabelGO+ incorporates the mean top sequence identity for dynamic weighting of AlignmentKNN, allowing the model to effectively leverage both sequence-based and language model-based features. A preliminary version of InterLabelGO participated in the Critical Assessment of Functional Annotation (CAFA) challenge [18] and achieved a strong performance, ranking 6th out of 1,625 participating teams. This demonstrates the efficacy of our approach in comparison to state-of-the-art methods.

## 2. MATERIALS AND METHODS

### 2.1 The Overall Framework of InterLabelGO+

The overall layout of the InterLabelGO+ is shown in **Figure 1**. The InterLabelGO framework begins with inputting the query protein sequence into the ESM2 [13] model, which generates a sequence embedding matrix from the last three hidden layers, with each layer producing a 3*×L×*2560 matrix where *L* represents the protein sequence length. Mean pooling is applied to the embedding vectors corresponding to each residue, resulting in compressed embedding vectors of length 2,560 from each hidden layer. These vectors represent embeddings that are further processed by three parallel multilayer perceptrons (MLPs). Each MLP is responsible for extracting evolutionary features from its corresponding layer, resulting in a 3 *×* 2560 matrix. These aggregated evolutionary data are then concatenated and processed by another MLP block. This layer’s purpose is to transform the ESM2-derived features into GO term probabilities. During the inference phase, a hierarchical post-processing approach is implemented, mandating that the probability of a parent term is at least equal to the maximum probability of its child terms, and raising the probabilities assigned to all non-leaf nodes in order to bring the predictions in line with this constraint.

**Fig. 1.**
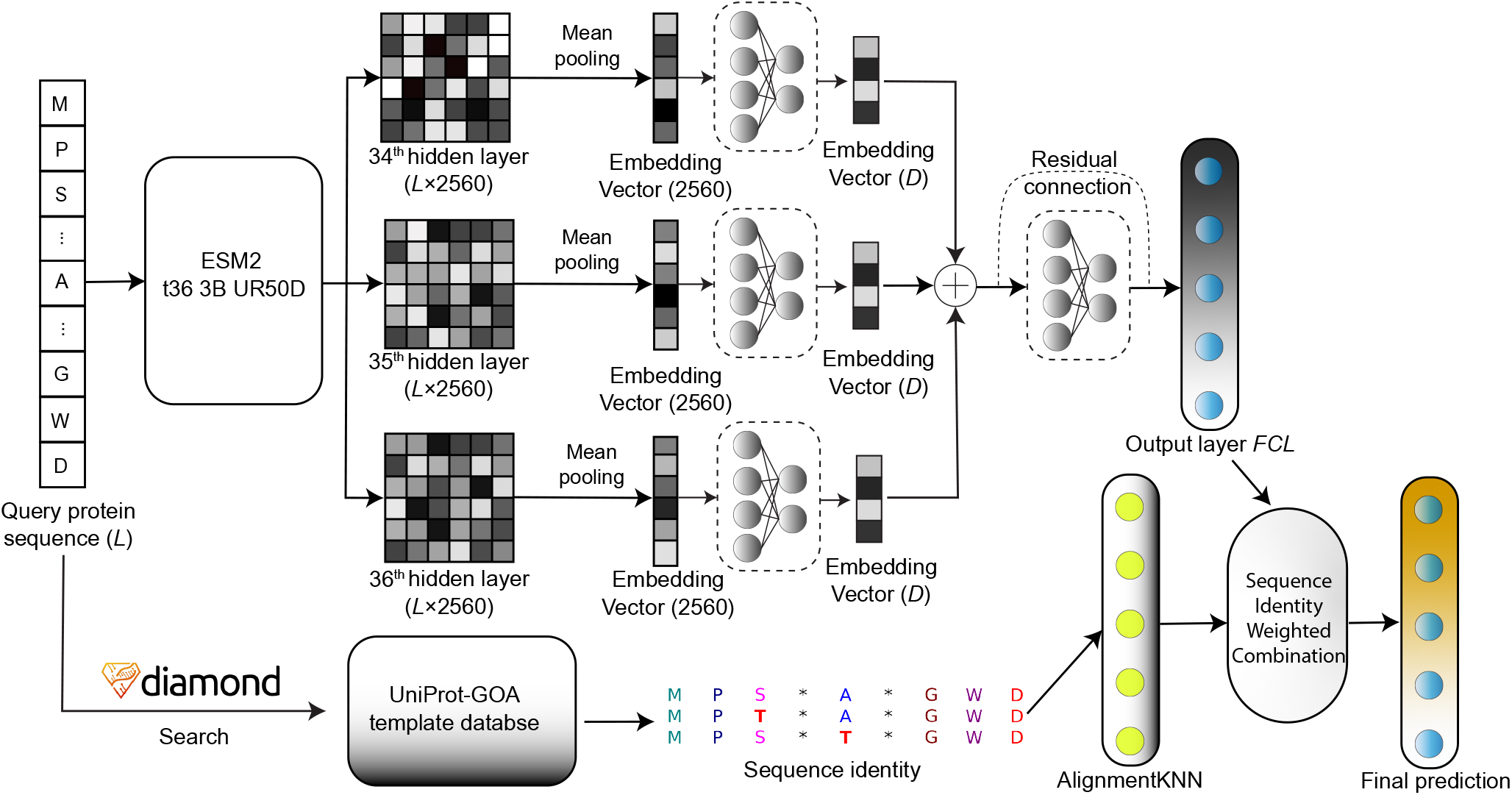
The workflow of InterLabelGO+.

In addition to the deep learning-based predictions from InterLabelGO, InterLabelGO+ also incorporates homology-based function transfer using AlignmentKNN, which is adapted from [16]. This approach involves searching the query sequence against a database of annotated template proteins using DIAMOND [1] and deriving prediction scores from normalized bitscores and sequence identity (detailed in Section 2.3). The final predictions of InterLabelGO+ are obtained by integrating the deep learning-based predictions with the homology-based predictions using a weighted combination approach (described in Section 2.4). This amalgamation of homology-based and deep-learning predictions provides the final output for protein function prediction.

### 2.2 Loss in Multi-label Classification

In the context of protein function prediction using GO annotations, we employ a composite loss function that addresses the challenges of class imbalance and captures label dependencies. The composite loss function consists of two components: an F1-score based loss that accounts for class imbalance and a rank-based loss that captures label dependencies. In the following subsections, we will define each component and then present the composite loss function.

#### F1-score based loss addressing class imbalance

A significant challenge in protein function prediction using GO annotations arises from the label imbalance among GO terms. This imbalance, where certain terms are over-represented and others are rare, can lead to suboptimal performance in standard loss functions, such as binary cross-entropy, as less frequent terms do not contribute sufficiently to the overall loss function.

To overcome this challenge, our approach incorporates a specialized F1 loss function, enhanced with the Information Accretion (IA) weight. Information Accretion, also known as Information Content, as introduced in [3], quantifies the additional information a term *q* contributes to an ontology annotation, assuming its parent nodes are already annotated. It is calculated as:

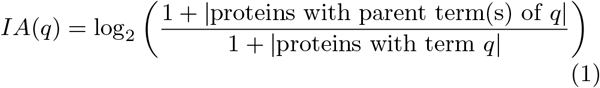

where |*X*| denotes the cardinality of set *X*.

The IA weight prioritizes GO terms with higher informational values, aiming to yield more informative predictions.

By factoring both precision and recall, the F1 loss function is inherently suitable for scenarios with label imbalance. The inclusion of the IA weight ensures that the function not only maintains a balance between precision and recall but also emphasizes the significance of more informative GO terms in the model’s learning process.

We employ two variants of the F1 loss: protein-centric and GO-centric. For the protein-centric F1 loss, precision and recall values are calculated for each protein in the batch, considering all its associated GO terms. In contrast, for the GO-centric F1 loss, precision and recall values are calculated for each GO term across all proteins in the batch.

For a single protein *i*, the precision and recall are calculated as follows:

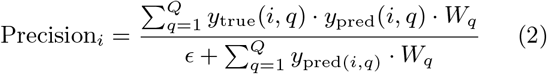

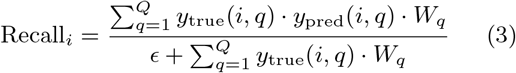

For a single GO term *q*, the precision and recall are calculated as follows:

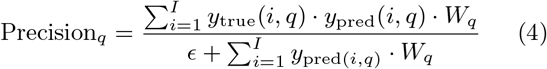

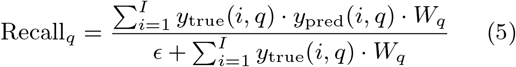

The F1 loss for both protein-centric and GO-centric variants is calculated as:

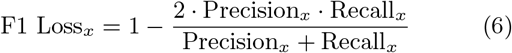

where *x* can be *i* (for protein-centric) or *q* (for GO-centric). In these equations, *I* is the total number of proteins in a batch, and *Q* is the total number of GO terms. *y*_true_(*i, q*) is the true label for protein *i* and GO term *q*, while *y*_pred_(*i, q*) is the predicted label for protein *i* and GO term *q. W*_*q*_ is the IA weight for GO term *q*, and *ϵ* is a small constant (1e-16) to avoid division by zero. For the protein-centric F1 loss (F1 Loss_*i*_), the mean precision and recall values are calculated over the protein dimension *i*. For the GO-centric F1 loss (F1 Loss_*q*_), the mean precision and recall values are calculated over the GO term dimension *q*.

#### Rank-based loss capturing label dependencies

The protein GO term prediction problem can be formulated as a hierarchical multi-label classification challenge. The unique challenge here is the structure of the GO term hierarchy, which form a large interconnected network, organized as three Directed Acyclic Graphs (DAGs) for the three GO aspects (MFO, BPO and CCO). This structure suggests that the prediction for one GO term may be influenced by other terms, acknowledging the inherent correlation and interdependence among them. In general, classification algorithms aim to determine the conditional probability *P* (*y* | *x*), capturing dependencies between input features x and target labels y. In multi-label classification, the complexity increases as dependencies may arise not just between features and target categories, but also among the categories themselves. Conventional methods, such as DeepGO [8], which typically utilize Binary Cross-Entropy (BCE) loss, often overlook the interdependencies between labels when treating each GO term as an independent binary classification problem. To better address these hierarchical and interdependent of GO terms, we adopt the Zero-bounded Log-sum-exp & Pairwise Rank-based (ZLPR) loss [12], shown in Equation 7.

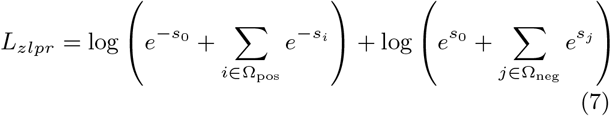

Here, *s*_*i*_ and *s*_*j*_ represent the logits from Output layer *FCL*, as shown in **Fig. 1**, for the i-th positive and j-th negative GO terms, respectively. Ω_*pos*_ and Ω_*neg*_ denote the sets of indices for the positive and negative GO terms. The ZLPR loss function introduces a pseudo term *s*_0_, set to 0 in our model, which acts as an intermediate threshold for distinguishing between positive and negative categories. *L*_*zlpr*_ represents the ZLPR loss for a single protein.

The ZLPR loss function effectively captures the dependencies among GO terms by leveraging information from their joint distribution. The formula distinctly differentiates between positive and negative categories using the log-sum-exp function and the pairwise ranking principle. By minimizing the ZLPR loss, the model aims to maximize the difference between the highest negative logit and the lowest positive logit, effectively ranking the positive categories above the negative ones through the intermediate term *s*_0_. This approach enables the model to consider the relationships between GO terms during the training process, rather than treating each GO term as an independent binary classification problem. Thus, it captures the complex correlations and dependencies among labels, which are crucial for accurate GO term prediction in the context of multi-label classification.

#### Composite Final Loss Function

To synthesize the strengths of the individual loss components, our final loss function is formulated as a multiplicative combination of the F1 loss calculated on a protein-centric basis, the F1 loss evaluated from a GO term-centric perspective, and the ZLPR loss. This composite approach ensures that the model optimally balances these different aspects during training, leading to enhanced prediction accuracy. The final loss function is defined as:

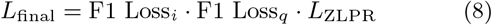

Here, F1 Loss_*i*_ represents the protein-centric F1 loss averaged across all proteins in the batch, F1 Loss_*q*_ denotes the GO term-centric F1 loss averaged for each GO term across all proteins, and *L*_ZLPR_ is the mean *L*_*zlpr*_ per protein, which captures the interdependencies among the GO terms. By integrating these components, the final loss function ensures a comprehensive and balanced learning process, taking into account both the individual and correlation-based characteristics of the protein function prediction task.

A detailed comparison of different loss functions and their combinations, along with their impact on the performance of InterLabelGO, is presented in Section 3.2.

### 2.3 Sequence homology-based AlignmentKNN

Homology-based function transfer is a foundational and extensively utilized method in predicting protein functions. This approach is based on the premise that proteins with similar sequences often exhibit similar functions. Within the AlignmentKNN framework, the sequence of a query protein is searched against the database of annotated template proteins using Diamond. The prediction score is derived from normalized bitscores and sequence identity as follows:

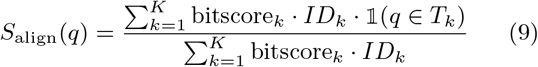

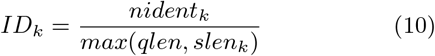

In this equation, *q* represents the *q*-th GO term. The terms *bitscore*_*k*_ and *ID*_*k*_ denote the bit-score and sequence identity of the k-th Diamond hit, respectively. The total number of hits considered, K, includes those hits which contain at least one GO term from the training set. *T*_*K*_ refers to the set of experimental annotations for the k-th target protein hit. The function is an indicator function that returns 1 if the GO term f is present in *T*_*K*_, and 0 otherwise.

The sequence identity *ID*_*k*_ for k-th protein hit is calculated in **Equation 10** where *nident* represents the number of identical amino acids found in the alignment between the query protein and the target. *qlen* and *slen* are denote the lengths of the query protein and the target protein, respectively.

### 2.4 Composite model of InterLabelGO+ using sequence identity

InterLabelGO+ is a weighted combination of AlignmentKNN and InterLabelGO. This approach is distinct from other methods, which typically rely on a linear combination of alignment scores and do not adequately capture complex relationships between scores, especially in cases where similar proteins are limited in number. InterLabelGO+ incorporates the mean top sequence identity for dynamic weighting, as detailed in **Equation 11 & 12**.

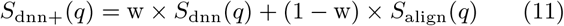

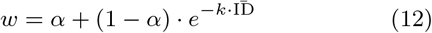

Here, *S*_dnn+_(*q*) represents the confidence score of InterLabelGO+ for the GO term *q*. The terms *S*_dnn_(*q*) and *S*_align_(*q*) denote the confidence scores for term *q* as predicted by InterLabelGO and AlignmentKNN, respectively. The weight parameter *w*, as defined in **Equation 12**, is calculated using the average of the top 5 sequence identities, represented by 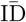 in **Equation (12)**. This average is combined with the minimum weight assigned by InterLabelGO, denoted as *α*. Additionally, the parameter *K* determines the rate at which the InterLabelGO weight varies with the average top sequence identities. The optimal values for *α* and *K* were established as 0.33 and 3, respectively, based on evaluations using the validation dataset.

### 2.5 Evaluation

To evaluate our predictions, we adopted the CAFA [18] challenge metrics, specifically the maximum weighted F-measure (*wF*_max_) and the minimum semantic distance (*S*_min_) [3]. In addition, we report the area under the weighted precision-recall curve(AUWPR), which is particularly relevant for datasets with significant label imbalance as it places greater emphasis on correctly predicting minority classes [4]. The AUWPR is calculated by plotting the weighted precision (wpr) against the weighted recall (wr) at various threshold settings and computing the area under the resulting curve where the weighted precision and recall are defined in **Equation 15**.

#### Weighted F-measure (*wF*_*max*_)

The metric *wF*_max_ represents the maximum of the protein-centric information accretion-weighted F-measure across all prediction thresholds *τ*. It is formulated as:

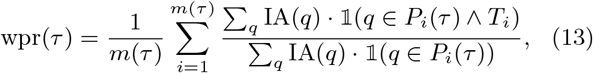

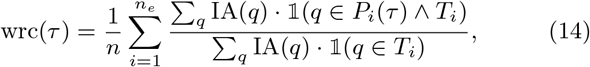

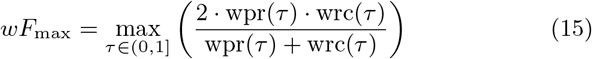

In this context, wpr(*τ*) and wrc(*τ*) denote the information accretion-weighted precision and recall at a given prediction score threshold *τ*. *P*_*i*_(*τ*) is the set of predicted terms for a protein *i* with a score equal to or higher than *τ*, and *T*_*i*_ represents the set of true terms for protein *i*. The number *m*(*τ*) indicates the count of sequences with at least one predicted score meeting or exceeding *τ*, and *n* is the total number of proteins in the benchmark. *q* represents the q-th GO term and IA(*q*) represents the information accretion of the q-th GO term as defined in the **Equation 1**.

#### Minimum Semantic Distance (*S*_*min*_)

The metric *S*_min_ calculates the semantic distance between actual and predicted annotations, considering the information accretion of classes. It is defined as:

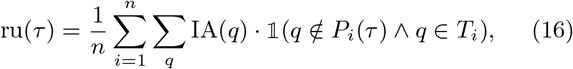

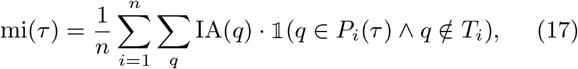

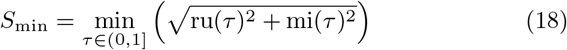

Here, ru(*τ*) represents the remaining uncertainty, and mi(*τ*) represents the missing information at threshold *τ*. For other terms, please refer to the **Section 2.5.1** for their definitions.

### 2.6 Datasets

We applied an approach closely paralleling the CAFA experiment for the benchmark datasets construction [18]. We downloaded all protein sequences from UniProtKB and GO term annotations from the Gene Ontology Annotation (GOA) database in November 2022, March 2023, and February 2024. We then filtered the proteins to include only those with at least one experimental annotation, identified by the following evidence codes: IDA, IPI, EXP, IGI, IMP, IEP, IC, or TAS. These annotations were subsequently propagated within the GO’s hierarchical structure to the root, using only the ‘is-a’ and ‘part-of’ relational criteria. The dataset was chronologically divided into training, validation, and testing subsets as follows: The training dataset comprised proteins annotated up to November 2022. The validation and testing datasets included no-knowledge and limited-knowledge proteins annotated between November 2022 and March 2023 and between March 2023 and February 2024, respectively. No-knowledge proteins are defined as those that lacked any experimental annotations prior to the start of the validation or testing period but gained at least one experimental annotation by the end of the corresponding period. In contrast, limited-knowledge proteins are characterized by having their first experimental annotations in the target GO aspects within the validation or test timeframe, while also possessing experimental annotations in at least one other domain before the start of the period. To optimize the training process and predictive accuracy, we only considered GO terms that met a minimum threshold of training instances. Specifically, we required at least 50 sequences for Biological Process (BPO) terms, and at least 10 sequences for both Molecular Function (MFO) and Cellular Component (CCO) terms. This criterion resulted in the inclusion of 2871 terms for the MFO, 5311 for the BPO, and 1626 for the CCO sub-ontologies, respectively. Detailed statistics on the benchmark datasets are shown on **Table 1**.

**Table 1.** Number of unique terms across different aspects for the training, validation, and test sets.

| Method | Training |  |  | Validation |  |  | Testing |  |  |
| --- | --- | --- | --- | --- | --- | --- | --- | --- | --- |
|  | BPO | CCO | MFO | BPO | CCO | MFO | BPO | CCO | MFO |
| Numbers of proteins | 92382 | 92920 | 78607 | 1146 | 699 | 563 | 5819 | 3440 | 2080 |
| Numbers of annotations | 3503437 | 1196126 | 669976 | 25252 | 7357 | 4081 | 169459 | 41592 | 19773 |
| Numbers of unique terms | 21321 | 2957 | 7224 | 3108 | 342 | 716 | 5781 | 689 | 1400 |

## 3. Results

### 3.1 Overall performance of InterLabelGO

To assess the performance of InterLabelGO, we compared it with four state-of-the-art composite models (ATGO+ [19], DeepGOPlus [7], TALE+ [2], and SPROF-GO [15]) on the test set (described in Table 1). We evaluated the models using three metrics: *wF*_*max*_, *S*_*min*_, and AUWPR. Additionally, we included an alignment-based method (AlignmentKNN) and a naïve baseline model for comparison, the naive method calculates the confidence score of a query being associated with a GO term by the frequency of that term in the training dataset. For the four third-party methods, we retrained the programs using the same dataset as InterLabelGO (detailed in Table 1) with their default training parameters and ran them on our test dataset under the default setting.

The results of the comparison are presented in Figure 2. InterLabelGO+ consistently achieves the best predictions for all three GO aspects when evaluated by all three metrics.

**Fig. 2.**
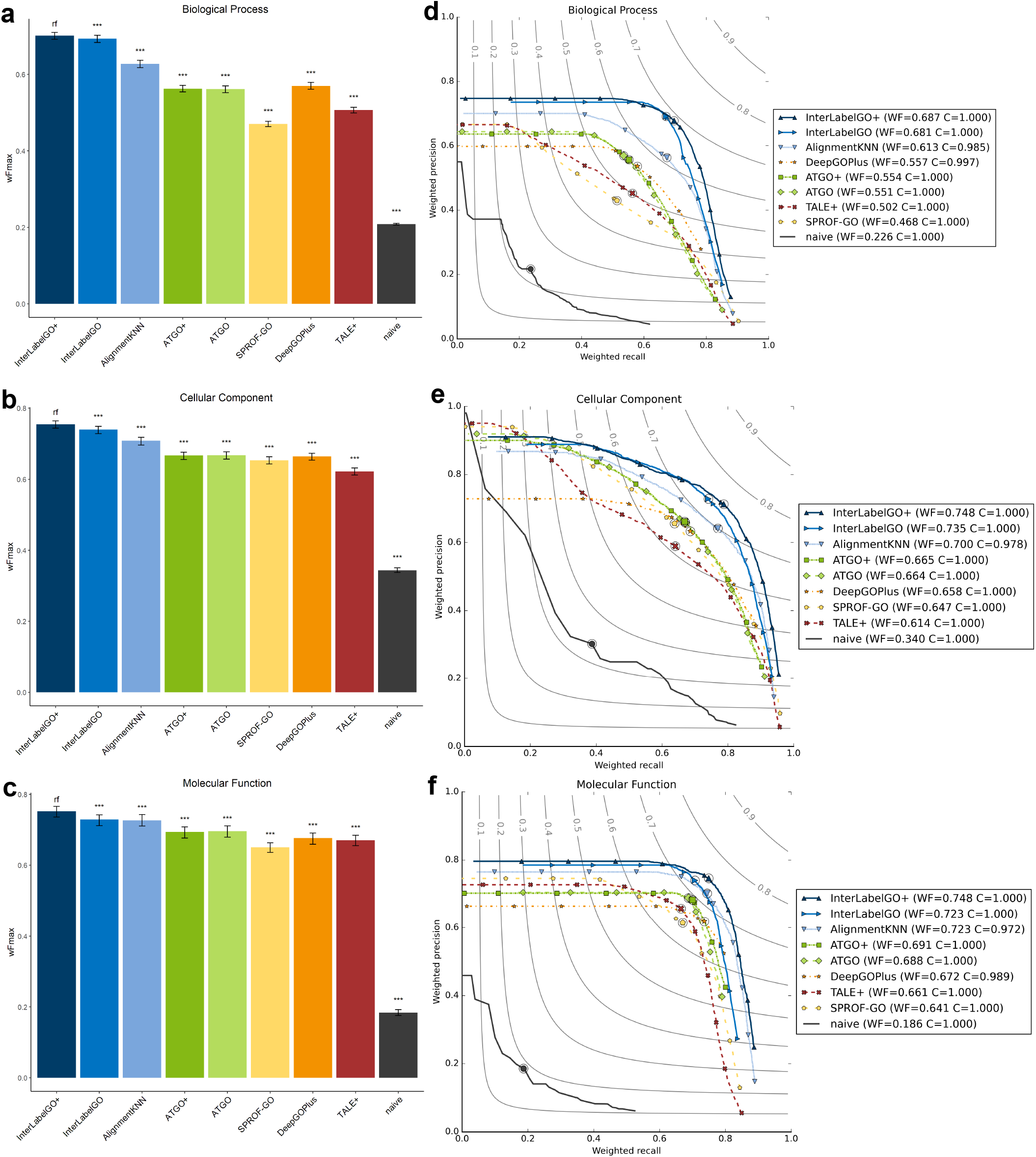
**Evaluation of Top-Performing Methods in Three Ontologies Using w***F*_**max**_. **Performance of different methods based on the UniprotGOA 202303-202402 release. Panels a–c display bar plots of the w***F*_**max**_ **for the different methods, with their 95% confidence intervals calculated via 9**,**999 bootstrap iterations on the benchmark dataset. Panels d–f illustrate the weighted precision-recall curves for these methods. Ideal performance, with w***F*_**max**_.**= 1, is represented at the top right corner of the plot, and the position of the maximum weighted F score on each curve is marked with a dot.The legend also indicates the coverage (denoted as C), which is the percentage of proteins in the benchmark predicted by each method, and the value of w***F*_**max**_.

It is noteworthy that the alignment-based method AlignmentKNN not only showed complementary performance to the neural network-based methods but also outperformed the other third-party deep learning-based methods in several cases. This observation suggests that as the number of experimental annotations grows in the database, the performance of alignment-based methods may also improve. The complementary nature of AlignmentKNN and InterLabelGO is particularly evident in the area under the weighted precision and recall curve (AUWPR) for the Molecular Function sub-ontology (Figure 2c). Although the *wF*_*max*_ scores for InterLabelGO and AlignmentKNN are similar, their precision-recall curves exhibit slightly different distributions, indicating that these methods capture different aspects of the protein function prediction problem and can complement each other.

The superior performance of our composite InterLabelGO+ method compared to the state-of-the-art methods highlights the effectiveness of our approach in capturing the complex relationships between GO terms and protein sequences. Furthermore, the complementary performance of the alignment-based method AlignmentKNN suggests that combining alignment-based and neural network-based approaches can lead to further improvements in protein function prediction.

### 3.2 Comparison of different loss functions and their combinations

To investigate whether the use of custom loss functions contributes to the success of InterLabelGO and to examine the effect of using different loss functions on the final performance, we trained different models with various loss functions and their combinations on the training dataset mentioned in Table 1. The loss functions considered in here include the Zero-bounded Log-sum-exp & Pairwise Rank-based (ZLPR) loss, Binary Cross-Entropy (BCE) loss, protein-centric F1 loss (PTF1), and GO-centric F1 loss (GOF1). The combinations of these loss functions were obtained by simply multiplying them together. The performance of these models was then evaluated using the test set described in Table 1 and presented in Table 2.

**Table 2.** Performance comparison of different loss functions and their combinations on the UniProt-GOA 20230316-20240209 release.

| Method | $wF_{max}$ | | | $S_{min}$ | | | AUWPRC | | |
| --- | --- | --- | --- | --- | --- | --- | --- | --- | --- |
|  | BPO | CCO | MFO | BPO | CCO | MFO | BPO | CCO | MFO |
| ZLPR_PTF1_GOF1 | 0.681 | **0.735** | 0.723 | 13.593 | 4.818 | 5.286 | 0.441 | 0.624 | 0.479 |
| ZLPR_GOF1 | **0.682** | 0.730 | **0.729** | **13.196** | 4.830 | **5.213** | 0.427 | 0.648 | 0.481 |
| ZLPR_PTF1 | 0.665 | 0.732 | 0.726 | 17.568 | 4.950 | 5.332 | 0.440 | 0.626 | 0.478 |
| ZLPR | 0.677 | 0.734 | 0.726 | 13.910 | **4.730** | 5.282 | 0.441 | 0.650 | 0.485 |
| BCE_ZLPR | 0.674 | 0.731 | 0.728 | 13.942 | 4.768 | 5.249 | 0.443 | 0.652 | 0.486 |
| BCE_PTF1_GOF1 | 0.661 | 0.717 | 0.684 | 13.914 | 5.142 | 6.115 | 0.464 | 0.620 | 0.500 |
| BCE_PTF1 | 0.647 | 0.713 | 0.695 | 14.690 | 5.158 | 5.873 | **0.466** | 0.619 | 0.499 |
| BCE_GOF1 | 0.642 | 0.706 | 0.702 | 14.037 | 5.183 | 5.735 | 0.434 | 0.674 | 0.505 |
| BCE | 0.627 | 0.701 | 0.694 | 14.785 | 5.258 | 5.855 | 0.446 | **0.675** | **0.515** |
| PTF1_GOF1 | 0.423 | 0.587 | 0.488 | 220.568 | 7.371 | 9.723 | 0.179 | 0.241 | 0.162 |
| PTF1 | 0.306 | 0.598 | 0.492 | 419.441 | 7.322 | 13.357 | 0.086 | 0.236 | 0.139 |
| GOF1 | 0.139 | 0.247 | 0.265 | 29.689 | 9.007 | 10.700 | 0.062 | 0.167 | 0.095 |

Best performance in bold. $wF_{max}$ and AUWPRC, highest; $S_{min}$ , lowest.

Our results show that adding PTF1 or GOF1 to BCE leads to improved performance, likely due to their ability to mitigate the label imbalance issue. The ZLPR loss itself provides an improvement over BCE, and further incorporating PTF1 and GOF1 enhances its performance, again by addressing the label imbalance problem.

Interestingly, the PTF1 and GOF1 losses, when used independently, perform poorly compared to other loss configurations including the classical BCE loss. This observation suggests that these losses, while effective in addressing label imbalance, may not be sufficient on their own to capture the complex relationships between GO terms and protein sequences. The combination of these losses with ZLPR or BCE appears to be crucial for achieving optimal performance.

Among the various loss combinations, the two best combinations are ZLPR GOF1 and ZLPR PTG1 GOF1 which perform similarly to each other. However, we ultimately selected ZLPR PTF1 GOF1 as the primary loss configuration for training InterLabelGO. This decision was based on its superior overall performance when combined with AlignmentKNN according to the validation dataset. The combination of ZLPR, PTF1, and GOF1 likely strikes a balance between capturing the pairwise ranking of GO terms, addressing label imbalance, and considering both protein-centric and GO-centric perspectives.

The results of our study underscore the importance of selecting an appropriate loss function or combination of loss functions for protein function prediction. The ZLPR loss, along with PTF1 and GOF1, demonstrates the effectiveness of addressing label imbalance and capturing the complex relationships between GO terms and protein sequences. The superior performance of ZLPR PTF1 GOF1 when combined with AlignmentKNN highlights its potential for robust and accurate predictions in real-world scenarios.

### 3.3 Advantage of hybrid model

To demonstrate the advantages of the hybrid approach employed by InterLabelGO+, we conducted a case study on three representative proteins from our test dataset, each with varying levels of sequence similarity to proteins in the GOA template database: Transcriptional regulatory protein dep1 (UniProt ID: Q9P7M1) from *Schizosaccharomyces pombe* (strain 972 / ATCC 24843), Glutamate receptor interacting protein 2 isoform X3 (UniProt ID: A0A8M6Z252) from *Danio rerio* (Zebrafish), and DEAD/H (Asp-Glu-Ala-Asp/His) box polypeptide 20 (UniProt ID: A6K3R6) from *Rattus norvegicus* (Rat). These proteins have 17, 25, and 32 experimentally annotated GO terms, respectively, under the BPO aspect, with all parent terms being propagated through ’is-a’ and ’part-of’ relationships. Table 3 presents a comparative analysis of the performance of InterLabelGO, InterLabelGO+, and seven other methods in the context of biological process (BPO) prediction for these proteins. The number enclosed in parentheses next to each protein UniProt ID represents the mean top 5 sequence similarity obtained by DIAMOND hits.

**Table 3.** **Performance comparison of InterLabelGO/InterLabelGO+ and seven other methods for BPO prediction on three representative proteins. Mean top 5 sequence similarity by DIAMOND is in parentheses. Best-performing method in each category is in bold**.

| Method | Q9P7M1 (N/A) |  |  |  | A0A8M6Z252 (0.61) |  |  |  | A6K3R6 (0.84) |  |  |  |
| --- | --- | --- | --- | --- | --- | --- | --- | --- | --- | --- | --- | --- |
|  | F1-score | FN | FP | TP | F1-score | FN | FP | TP | F1-score | FN | FP | TP |
| InterLabelGO+ | **0.774** | 5 | 2 | 12 | **0.939** | **2** | 1 | **23** | **0.955** | **0** | 3 | **32** |
| InterLabelGO | 0.741 | 7 | **0** | 10 | 0.913 | 4 | **0** | 21 | 0.731 | 13 | **1** | 19 |
| AlignmentKNN | 0.000 | 17 | 0 | 0 | 0.177 | 18 | 47 | 7 | 0.853 | 0 | 11 | 32 |
| ATGO+ | 0.516 | 9 | 6 | 8 | 0.324 | 19 | 6 | 6 | 0.356 | 24 | 5 | 8 |
| ATGO | 0.562 | 8 | 6 | 9 | 0.324 | 19 | 6 | 6 | 0.391 | 23 | 5 | 9 |
| TALE+ | 0.364 | 9 | 19 | 8 | 0.250 | 19 | 17 | 6 | 0.831 | 0 | 13 | 32 |
| SPROF-GO | 0.395 | **2** | 44 | **15** | 0.290 | 16 | 28 | 9 | 0.487 | 13 | 27 | 19 |
| DeepGOPlus | 0.000 | 17 | 0 | 0 | 0.389 | 18 | 4 | 7 | 0.615 | 0 | 40 | 32 |
| naive | 0.250 | 13 | 11 | 4 | 0.300 | 19 | 9 | 6 | 0.170 | 28 | 11 | 4 |

For each method, the prediction score threshold used in the case study corresponds to the value that achieved the best *F*_*max*_ on the entire test dataset. The table highlights the top-performing method in each category, marked with bold font. Figure 3 displays the DAG of GO terms for the protein A0A8M6Z252. The number in parentheses next to each GO term represents its Information Accretion (IA) value. The methods that correctly predicted each GO term are listed below the corresponding term.

**Fig. 3.**
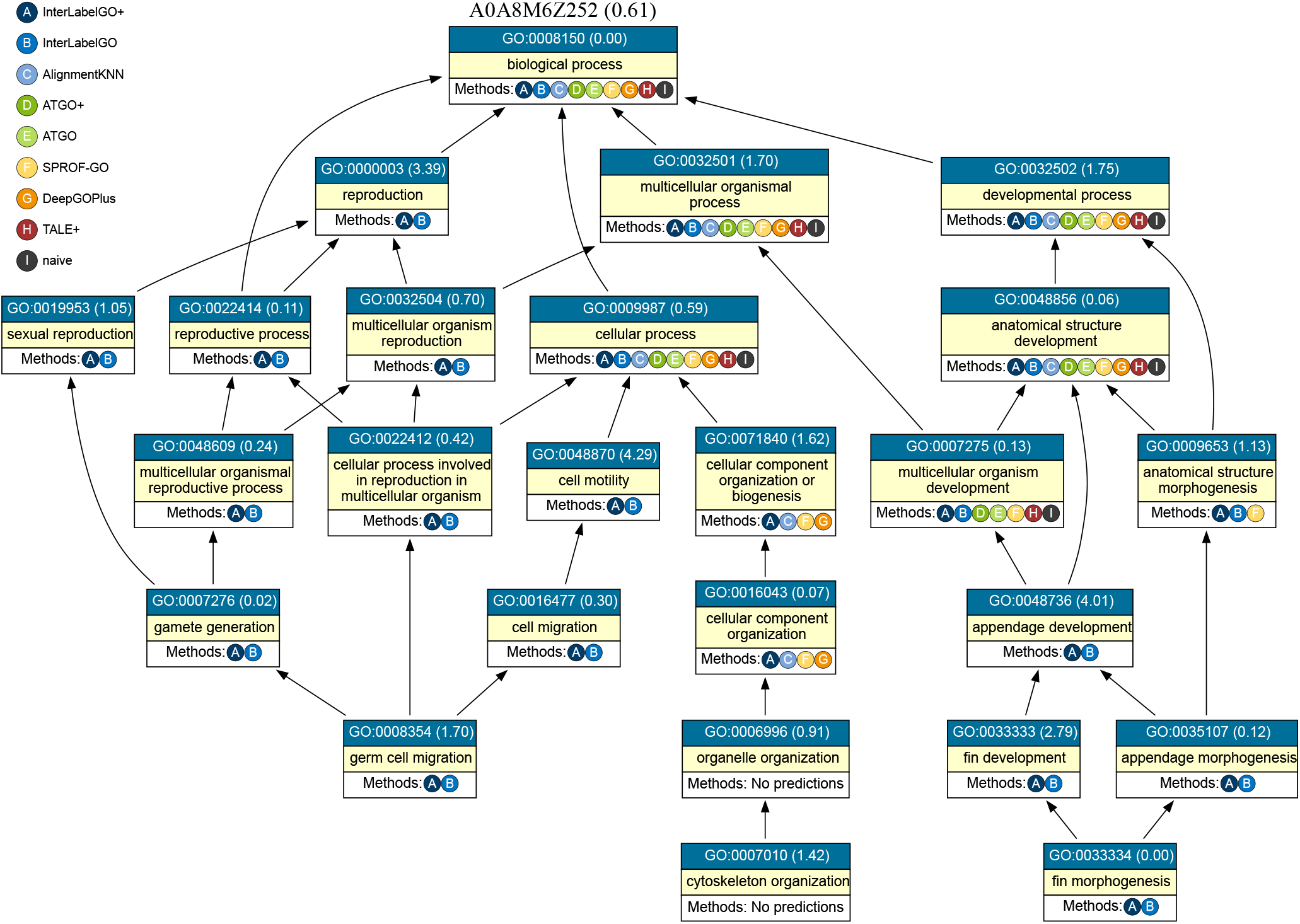
**Directed acyclic graph showing the biological process GO terms for the representative example A0A8M6Z252. All methods that correctly predicted each GO term are listed below the corresponding term. The number in parentheses next to each GO term represents its Information Accretion (IA) value**.

The case study results reveal the strengths of InterLabelGO and InterLabelGO+ in capturing complex relationships between GO terms and predicting groups of related terms across various sequence similarity scenarios.

For the challenging protein Q9P7M1, which lacks significant hits in the GOA template database, InterLabelGO demonstrates its suitability by maintaining relatively good performance compared to all other methods. When considering composite methods that combine alignment-based and deep learning approaches, InterLabelGO+ still performs the best. Unlike ATGO+ and ATGO, where the combination of deep learning and alignment-based methods can lead to decreased performance if the alignment-based component fails to provide reliable predictions, the dynamic weighting scheme in InterLabelGO+ helps mitigate the negative impact on the overall results.

In the case of A0A8M6Z252, where the alignment-based method has hits but exhibits a high level of false positives, InterLabelGO+ showcases the complementary nature of InterLabelGO and AlignmentKNN predictions. This combination effectively filters out most of the false positives from AlignmentKNN while incorporating true positives that are not present in InterLabelGO, further improving the overall score.

For A6K3R6, where the GOA template database contains very similar proteins, AlignmentKNN performs exceptionally well, surpassing all other methods, including InterLabelGO and the composite method TALE+. The combination of alignment-based predictions and InterLabelGO in InterLabelGO+ maintains the high true positive rate from AlignmentKNN while reducing false positives, again demonstrating the complementary nature of these approaches and the importance of the hybrid approach.

A closer examination of A0A8M6Z252 (Figure 3) reveals that InterLabelGO is capable of predicting a large group of functions that all other methods failed to identify, including GO terms with high Information Accretion (IA) values such as GO:0000003 (3.39), GO:0048870 (4.29), and GO:0048736 (4.01). The incorporation of IA and label correlation in the loss function enables InterLabelGO to capture intricate relationships between GO terms, allowing it to predict a significantly larger number of terms compared to other methods.

However, the case study also highlights a few instances where InterLabelGO missed some terms, such as GO:0071840 and GO:0016043, which were successfully predicted by the combination of AlignmentKNN and InterLabelGO+. This observation underscores the importance of integrating alignment-based and deep learning-based methods to achieve comprehensive and accurate GO term predictions.

The case study provides valuable insights into the performance of InterLabelGO and InterLabelGO+ in real-world scenarios, emphasizing their ability to capture complex GO term relationships and the benefits of combining different approaches for optimal results across various levels of sequence similarity.

## 4. DISCUSSION

InterLabelGO is a novel alignment-free deep learning-based approach that accurately predicts protein functions from sequence information alone. Our model addresses several challenges in protein function prediction, including label imbalance and label dependencies, by incorporating a composite loss function that combines a GO term-aware weighted F1 loss and a pairwise ranking-based loss. This enables InterLabelGO to capture complex functional relationships and mitigate the impact of skewed functional annotation distributions. Furthermore, InterLabelGO integrates sequence homology-based predictions through a dynamic weighting scheme, leveraging complementary information to enhance its predictive performance. The superior performance of InterLabelGO compared to state-of-the-art methods across various functional categories and evaluation metrics demonstrates its effectiveness in unraveling the functional landscape of proteins. Moreover, the strong performance of InterLabelGO in the CAFA challenge highlights its potential for real-world applications and its ability to compete with top-performing methods in the field.

In the future, we plan to explore additional features and architectures to further improve the predictive performance of InterLabelGO+. One potential approach for improvement is to incorporate attention mechanisms within the InterLabelGO framework. Attention mechanisms have been successfully employed in various natural language processing tasks and have recently shown promise in protein function prediction, as demonstrated by SPROF-GO [15]. By integrating attention mechanisms, we aim to enable InterLabelGO+ to focus on the most informative regions within protein sequences and provide insights into the key patterns contributing to specific functions. This could not only enhance the model’s predictive performance but also improve its interpretability. Furthermore, we intend to extend the deep learning model of InterLabelGO to handle multi-modal data, such as protein-protein interaction networks and literature-derived features. Protein-protein interaction data can provide valuable information about the functional associations between proteins, while literature-derived features can capture relevant functional information from the vast body of scientific literature. Techniques like BioBERT [10] and the approach used in DeepText2GO [14] could be leveraged to extract meaningful features from the literature. By incorporating these additional data sources, we aim to provide InterLabelGO+ with a more comprehensive understanding of protein functions and further improve its predictive capabilities.

